# Protein level variability determines phenotypic heterogeneity in proteotoxic stress response

**DOI:** 10.1101/646653

**Authors:** Marie Guilbert, François Anquez, Alexandra Pruvost, Quentin Thommen, Emmanuel Courtade

**Author notes:** Corresponding Author: Quentin Thommen.

## Abstract

Cell-to-cell variability in stress response is a bottleneck for the construction of accurate and predictive models that could guide clinical diagnosis and treatment of diseases as for instance cancers. Indeed such phenotypic heterogeneity can lead to fractional killing and persistence of a subpopulation of cells resistant to a given treatment. The heat shock response network plays a major role in protecting the proteome against several types of injuries. We combine high-throughput measurements and mathematical modeling to unveil the molecular origin of the phenotypic variability in the heat shock response network. Although the mean response coincides with known biochemical measurements, we found a surprisingly broad diversity in single cell dynamics with a continuum of response amplitudes and temporal shapes for several stimuli strengths. We theoretically predict that the broad phenotypic heterogeneity is due to network ultrasensitivity together with variations in the expression level of chaperons controlled by heat shock factor 1. We experimentally confirm this prediction by mapping the response amplitude to concentrations chaperons and heat shock factor 1 expression level.

## Introduction

Resistance of a population subfraction to a cancer treatment (chemotherapy for instance) limits the effectiveness of this treatment (LeBlanc *et al*, 2002) and is named cellular response heterogeneity. Obviously, extracellular environment variations or genetic alterations induce cellular heterogeneity in treatment response. But several massive single-cell experimental results (Albeck *et al*, 2008; Feinerman *et al*, 2008; Gascoigne and Taylor, 2008; Orth *et al*, 2008; Irish *et al*, 2004; Cohen *et al*, 2008; Geva-Zatorsky *et al*, 2006) reveal that a significant phenotypic heterogeneity persists even for monoclonal cell lines and in uniform environment. The discovery of the underlying molecular mechanisms leading to variability in response to treatment and their potential control is a major issue for cancer therapy research (Niepel *et al*, 2009; Almendro *et al*, 2013).

In a clonal cell line, intracellular biochemical fluctuations create a cell population with the same genome but with various proteomes (Kærn *et al*, 2005; Sigal *et al*, 2006). Cell-to-cell variability can arise from such intracellular biochemical fluctuations and is called Non-Genetic Heterogeneity (NGH) (Huang, 2009). NGH plays a functional role in surviving unpredictable environmental changes (Kærn *et al*, 2005; Acar *et al*, 2008; Pfeuty and Thommen, 2016), and it has been identified in anticancerous treatment as a inducer of fractional killing (Spencer *et al*, 2009; Flusberg and Sorger, 2015; Roux *et al*, 2015). Accurate clinical models including NGH are still to be built in order to guide diagnosis and treatment of diseases (Bertaux *et al*, 2014). Indeed the precise knowledge of the molecular network is not enough to predict the response of a cell population to a given treatment. Such models would also require the identification of the key molecular players (Behar *et al*, 2013; Reyes and Lahav, 2018) and a detailed study of NGH (Loewer and Lahav, 2011; Altschuler and Wu, 2010).

A general feature of cellular stress response networks is the response-to-stimuli ultrasentivity: the response increases slowly at low stimuli value and sharply increases to high response once a given stimulus threshold is reached. Ultrasensitivity is well known to arise from networks having either a positive cooperativity, multistep processes or protein sequestration (Goldbeter and Koshland, 1984; Buchler and Cross, 2009). Such architectures tend to shrink Cell-to-Cell Response Variability (CCRV) due to NGH for low or moderate stress strengths. In contrast, the response-to-stimuli ultrasensitivity tends to broaden CCRV once the stimulus approaches the threshold value. Stress response ultrasensitive networks are thus appropriate to study the cellular heterogeneity arising from NGH because ultrasensitivity acts as a heterogeneity amplifier.

The Heat Shock Response Network (HSRN) in the cytosol, together with the unfolded protein response in the endoplasmic reticulum, is essential for maintaining the proteome integrity (Morimoto, 2012). HSRN displays ultrasensitivity due to protein sequestration mechanism (Buchler and Cross, 2009; Sivéry *et al*, 2016). Several proteotoxic stresses, such as oxidation or heat, trigger HSRN which induces the transcription of Heat Shock Proteins (HSPs) *via* activation of Heat Shock Factor 1 (HSF1) transcription factor. HSPs act as molecular chaperones to maintain proteostasis (Jolly and Morimoto, 2000).

At the single cell level, HSF1 forms dynamic structures named Nuclear Stress Bodies (nSBs) (Biamonti and Vourc’h, 2010). In the present study, we first show that nSBs can be used to quantify HSRN activation at the single cell level. We then use high-throughput time-lapse microscopy with a precise control of hyperthermia temporal profile (41°C-43°C) to monitor nSBs dynamics in a monoclonal population. From computational image analysis large data sets of quantitative single cell nSBs temporal dynamics have been constructed several hours after heat shock. These data allow us to shed light on an unexpected broad range of response to a given stimulus. We then address the molecular underpinnings of such CCRV in HSRN dynamics. Response variability is investigated by both statistical analysis of data and network parameters sensitivity analysis of a data-driven mathematical description of the HSRN. Using computational prediction and experimental characterization of single cells, we finally identify NGH to play a crucial role in CCRV by modulating HSF1 and HSPs concentration across the cell population.

## RESULTS

### Monitoring HSF1 activation of individual HeLa cells

Under normal conditions the molecular chaperon *heat shock protein 70 kDa* (HSP70) sequesters HSF1. Under stressful conditions HSF1 is unbound from HSP70 (Abravaya *et al*, 1992; Kline and Morimoto, 1997). Free HSF1 can form homotrimers (Sarge *et al*, 1993; Cotto *et al*, 1996) and can bind on specific region of DNA named *Heat Shock Elements* (HSE). HSF1 bound to HSE promotes the transcription of the wide family of HSP proteins that includes HSP70 and others (Mosser *et al*, 1988; Baler *et al*, 1993; Holmberg *et al*, 2002; Cotto *et al*, 1997; Boulon *et al*, 2010). In human and primate cells, free HSF1 also form *Nuclear Stress Bodies* (nSBs also named as *HSF1 foci*) which are reversible macromolecular complexes made of (among other macromolecules) HSF1 trimers bound to heterochromatin regions without HSE (Biamonti and Vourc’h, 2010). If HSF1:eGFP binding to HSE provides an insufficient fluorescent signal to be efficiently monitored using conventional fluorescence microscopy a single nSB does (Cotto *et al*, 1997; Biamonti and Vourc’h, 2010). nSBs are formed within seconds under stressful conditions (Biamonti and Vourc’h, 2010) and form foci in the cell nucleus that can be observed under a conventional fluorescence microscope (Fig. 1 A and Cotto *et al*, 1997). The quantity of HSF1:eGFP within foci can be measured over statistically significant cell population by the use of an automated images analysis. We define *F* as the fraction of HSF1:eGFP fluorescence signal located in foci to measure HSR activation for an individual cell. *F* is a ratiometric measurement proportional to the fraction of HSF1 free from HSP70. *F* provides a readout of HSF1 activation in individual cells (Fig. 1A).

**Figure 1:**
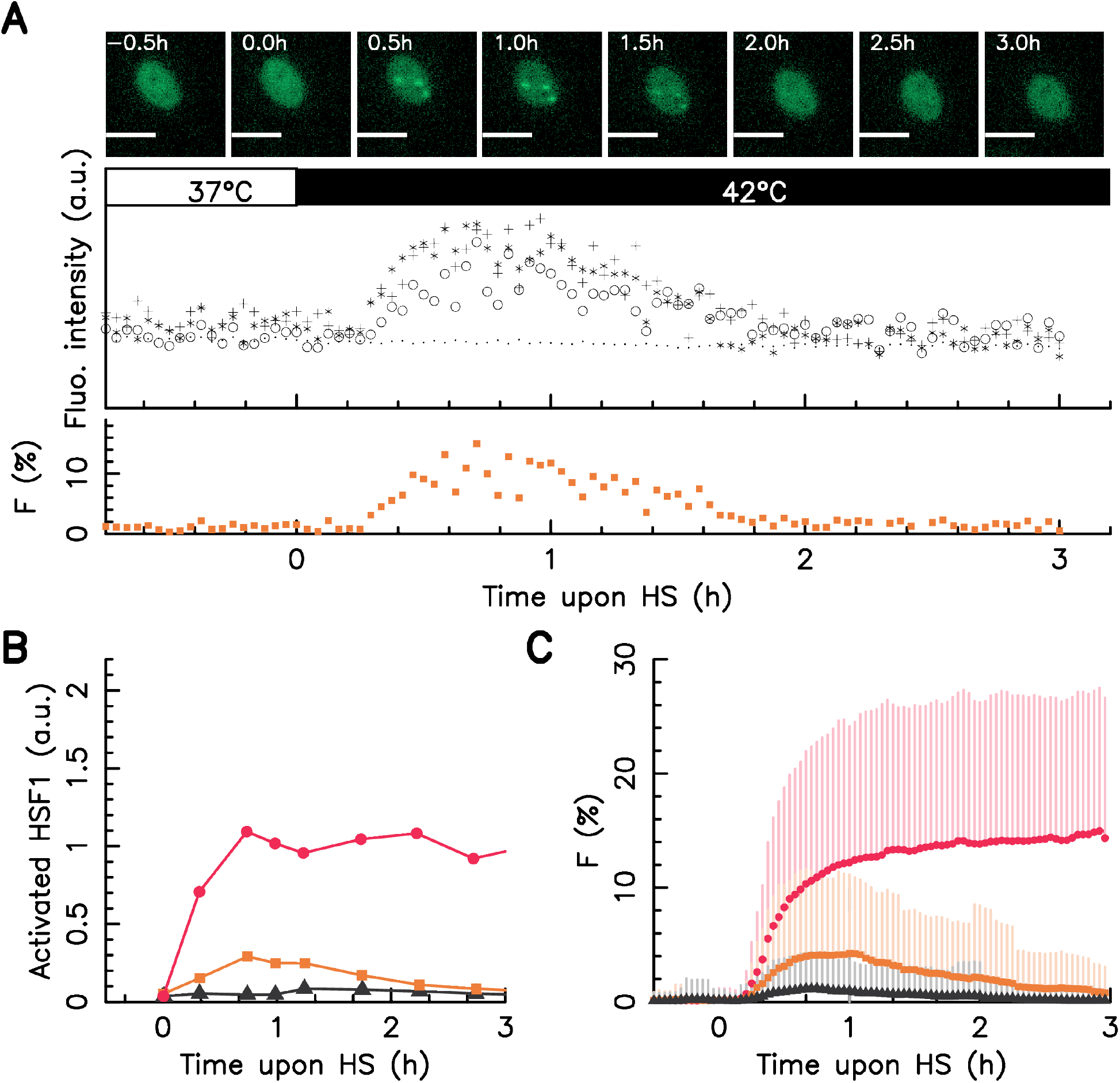
Screening of foci dynamics in individual HeLa cells. A – upper panel: snapshots of single HeLa HSF1:eGFP cell over time upon a 42°C heat stress, in the images, the white scale bars correspond to 10 *μ*m; middle panel: dynamics of the fluorescence intensity measured in the center of the three visible foci (crosses, circles and starts) and average fluorescence level over the entire cell nucleus (dots); bottom panel: dynamics of the fraction of HSF1:eGFP fluorescence within foci *F*. B – Dynamics of activated HSF1 as measured by Abravaya *et al*, 1992 upon a 41°C (black) 42°C (orange), and 43°C (red) heat stress by run on assay C – Dynamics of the faction of HSF1:eGFP fluorescence (*F*) in nSBs monitored over time average over the whole cell populations upon a 41°C (black) 42°C (orange), and 43°C (red) heat stress for a large cell colony, dots stand for average values with on side error bars (standard deviation). In all cases, time zero coincides with the stress onset.

In order to compare our single cell method with conventional biochemistry measurements we first study the dynamics of *F* averaged over the whole cell population upon a temperature rising up from 37°C to 41, 42, or 43°C. As shown by Abravaya *et al*. (Abravaya *et al*, 1992) dynamics of HSF1 bound to HSE at these three temperatures deliver the big pictures of the HSR dynamics upon heat stress. Indeed the dynamics of activated HSF1 at 42°C is drastically different from the one at 43°C: at the former temperature activated HSF1 exhibits pseudo adaptation kinetics while at the later temperature HSR activation persists (Fig. 1-B). The time evolution of *F* average over the whole population is in very good agreement with biochemical measurements for all three temperatures (Fig. 1 C). The genetic modification resulting from the HSF1:eGFP insertion does not impact the *F* kinetic, as revealed by time point immunofluorescence staining measurement in wild-type HeLa (HeLa WT) cell line (see Fig. SI 1 A of the supporting information). We conclude that HSF1 foci dynamics as a valid reporter of the HSRN activation upon heat shock.

### High-troughput screening of HSRN reveals broad cell-to-cell variability

Averaging over the cell population gives at first glance a misconception: a population of cells all having similar foci intensity whose brightness increases with stress amplitude. Examining the time traces of individual cells with identical genome and exposed to the exact same stimulus reveals that this picture is not correct. One easily distinguishes a broad cellular heterogeneity both in foci intensity and dynamics (Fig. 2 A-C). These experimental results were confirmed using a second monoclonal cell line (see Fig. SI 3 of the supporting information). Besides no spatial dependency was found for the amplitude neither the shape of the response. This confirms that heterogeneity is not due to a spatial distribution of the temperature across the sample nor any other imaging artifact. Although HSF1:eGFP expression level varies significantly from cell–to–cell the total HSF1:eGFP amount in a single cell does not vary during the experiment and then does not impact the foci dynamic (see Fig. SI 2 A-C of the supporting information).

**Figure 2:**
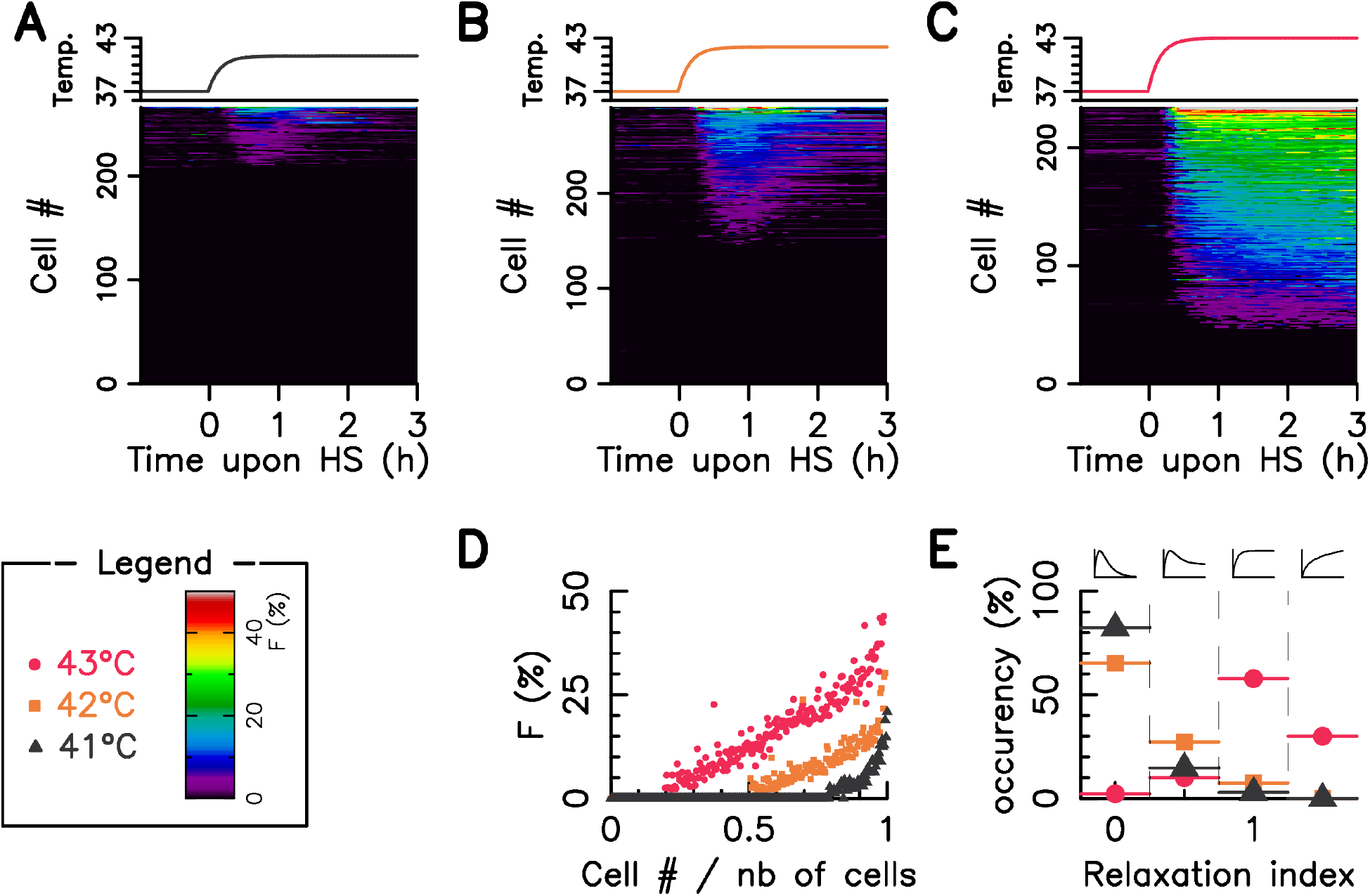
Cell-to-cell variability in heat shock response. The faction of HSF1:eGFP fluorescence in nSBs (*F*) is monitored over time in single cell upon a 41°C 42°C and 43°C heat stress, time zero coincides with the stress onset. A-C Cell temperature time profile (upper panel) and *F* as a function of time in a single cell (lower panel); in the color image, each horizontal line corresponds to a single cell, a color code indicates the *F* value measured at a given time. D – Distribution of *F* across the cell population for several heat shock temperatures one hour after the stress onset (cell ranking is similar to D-F). E – Statistical distribution of the relaxation index defined as the ratio of the foci intensity measured in a given cell at three hours after the stress onset to the one measured one hour after the stress onset. The legend box defines the used color code.

If we now focus on the response at a given time (one hour after the stress onset) we observe a significant fraction of cells that do not display detectable foci (78.4 % at 41°C, 50.5 % at 42°C, and 19.7 % at 43°C) while the responding subset displays a wide distribution of free HSF1 with *F* ranging from 0 to nearly 0.5 (Fig. 2-D). These numbers suggest that the rise of response amplitude with increasing temperature observed at the population level (Fig. 1 B and C) is at least partially due to an increase of the fraction of responding cells rather than solely due to an absolute increase in free HSF1 fraction per cell. Similar results are obtained with wild-type HeLa cell line (see Fig. SI 1 B of the supporting information).

The temporal shape of the response also varies across the cell population. We observe cells exhibiting complete relaxation and cells with *F* monotonously increasing. To capture the heterogeneity of HSF1 activation dynamics we define a relaxation index *η* as the ratio of the response at 1h to the one at 3h post heat shock (Fig. 2-E). *η* = 0 indicates a near perfect adaptation; *η* = 1 translates into a plateau; and *η* > 1 is a sign of continuously increasing activation. It is worth noting that some cells exposed to step temperature increase at 42°C show dynamics comparable to the population average at 43°C and *vice versa*. Indeed nearly 10 % of the responding cells have a relaxation index *η* > 1 for 42°C heat shock whereas 10 % of the responding cells shows a relaxation index *η* < 0.5 at 43°C. Finally we note that the temporal shape of the response is positively correlated with the *F* value one hour after the stress onset: the brighter is the foci the less pronounced is the relaxation (see Fig. SI 2 D-F of the supporting information).

### Variation of protein basal expression level can induce heterogeneous cellular response

One surprising feature of our single cell dataset is the apparently continuously varying behavior across the cell population (Fig. 2 A-C). Our attempts to apply statistical clustering methods to each dataset could not converge towards a finite number of phenotypes. A situation with only two clusters corresponding to the responding cells, on the one hand, and undetectable responses, on the other hand, is not satisfying as it would hide heterogeneity in the former class. We concluded that the variety of kinetic traces could not be captured by a discrete set of typical behavior. At the network level HSR is characterized by two competitive sequestration mechanisms. The output of such motif is known to be highly sensitive to protein concentration (Goldbeter and Koshland, 1984; Buchler and Cross, 2009). To explain the observed heterogeneity we hypothesized that a variation in basal protein expression level across cell population could lead to significant differences in cellular responses. Indeed protein expression levels vary from one cell to another even in a monoclonal cell line. This can be due to the stochastic expression of the gene (Sigal *et al*, 2006) or asymmetric cell division (Neumüller and Knoblich, 2009). To assess this possibility and gain understanding on the origin of CCRV we derived a coarse-grained mathematical model of the HSR network.

In a minimal description HSRN involves three different species: *(i)* MisFolded proteins (MFP) that are heat-induced and *(ii)* HSP which helps to refold MFP and *(iii)* HSF1 that promotes transcription of HSP. The dynamics of the network is mainly regulated by two complexes that both involve the chaperon HSP (Sivéry *et al*, 2016): HSPs titrate the MFP on the one hand and its own transcription factor (HSF1) on the other. Our model accounts for the temporal evolution of copy number of four molecular species (MFP; *HSP* mRNA; HSF1; HSP). The model is using ordinary differential equations where the fast dynamics of molecular complex assembly and disassembly are adiabatically eliminated (see Material and Methods for details). We also account for the measured temperature rise time of the incubator. The coarse-grained model is accurate enough to quantitatively describe the foci dynamics (see Fig. SI 4 of the supplementary information for comparison to experimental data and parameters estimation).

Using the above describe mathematical model, we show that reducing or increasing by only two fold the basal HSP concentration is sufficient to qualitatively mimic the dynamics of *F* (Fig. 3 A-C and D-F compared to Fig. 2 A-C). In our mathematical framework the response heterogeneity is captured as a consequence of protein copy number variability: the more is the HSP number, the less is the foci intensity (*F_th_*) and the lower is the relaxation index (*η_th_*). Moreover the population level observations are also predicted: *(i)* both *F_t_h* and *η_th_* increase with temperature and *(ii)* the temporal shape the foci intensity display more relaxation (*η_th_* < 0.5) at 41°C and 42°C than 43°C; a plateau or a slow increase (*η* ⩾ 1) is observed mostly at 43°C. Similar *in silico* results were obtained by varying HSF1 concentration (see Fig. SI 5). We conclude that variations of both HSF1 and HSP expression levels could lead to the experimentally observed CCRV.

**Figure 3:**
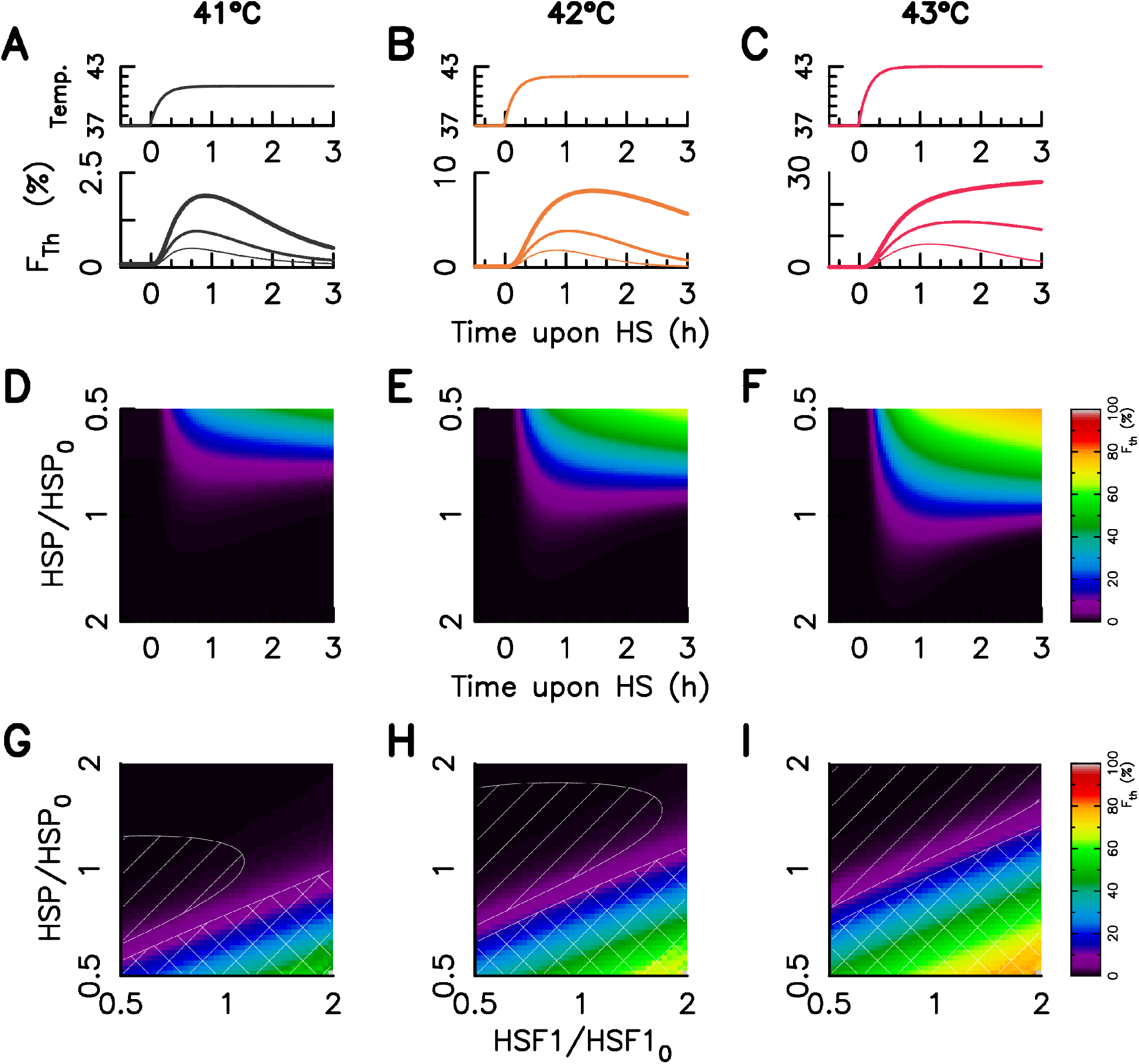
Mathematetical Modeling of heat shock response variability. A–C – Temperature time profile (upper panel) and three examples of predicted foci dynamics for a given HSF1 and three various HSP level of expression, the thicker the line, the more the HSP level of expression (lower panel); D–F – Foci dynamics dependency on the HSP level of expression in the case of a 41°C (D), 42°C (E), and 43°C (F) heat stress. G-I – Foci intensity one hour after the stress onset for varying HSP and HSF1 copy number. HSP and HSF1 levels are expressed in fold change of the concentration for the model fitted on the population averaged data sets(HSF1_0_ and HSP_0_). The relaxation index value is below 0.5 in the linear hatched area and above 1 in the crosshatched area. The stress intensity is indicated on the top of all columns. The used color code is similar to Fig 1 and is indicated in the legend box.

One of the major advantage of our model is that it provides an explicit analytical expression for the foci intensity *F_th_*(*t*) at any time *t* after the stress onset (Eq. 8). *F_th_*(*t*) depends on the concentration of the three main molecular actors, namely HSP, HSF1 and MFP. Mapping *F_th_* as a function of HSP and HSF1 concentrations reveals iso-response lines (Fig. 3 G-I). Such a mapping can be used to test the theoretical prediction experimentally.

### HSP72 and HSC70 expression level impact cellular response and lead to cell-to-cell heterogeneity of the HSR

Modeling results suggest that both HSP and HSF1 level variations may induce the observed cell-to-cell variability in response to heat stress. Therefore in order to test our theoretical predictions we use immunolabelling and fluorescence microscopy to measure simultaneously HSP and HSF1 concentrations together with the response *F* at the single cell level. Our model assumes a generic HSP while the HSP family is wide an comprise several variants with specific roles (Whitley *et al*, 1999). However only the HSP70 subfamily appears to play a significant role in HSF1 titration and consequently in its activation (Shi *et al*, 1998). We thus consider only two members of the HSP family: HSP72 and HSC70. Both proteins play a similar role in sequestrating HSF1 and the refolding of MFP (Gething and Sambrook, 1992), but HSP72 is a stress inducible protein (the transcription rate of *hsp72* mRNA increases with the free concentration of HSF1) whereas HSC70 is not stress induced and constitutively expressed (Tavaria *et al*, 1996).

In a first step we estimated HSF1, HSP72 and HSC70 concentration in individual cells from single immunolabeling in both HeLa Wild Type and in HeLa–HSF1:GFP cell lines. Experimental data were compared for two thermal conditions: without heat shock and one hour after exposure to a temperature step-up from 37°C to 43°C (Fig. 4 A-C). As HSF1 is located in the cell nucleus (see (Mercier *et al*, 1999 and Fig. 1 A) we focus on nuclear concentration for all three proteins. We used Hoechst staining of cellular DNA to allow automated cell segmentation of the cell nucleus. All three protein concentrations at both 37°C and 43°C are well fitted by a log-normal distribution (see Fig. SI 6 of the supporting information). As expected only HSP72 exhibits a shift of the distribution upon heat stress (Fig. 4 B). The HSF1:eGFP insertion induces an overexpresion of both HSF1 and HSP72 (1.46 for HSF1 and 1.65 for HSP72) but has no effect on HSC70. HSF1:eGFP insertion also induces a broadening of the HSF1 and HSP72 distribution especially toward higher values for both species.

**Figure 4:**
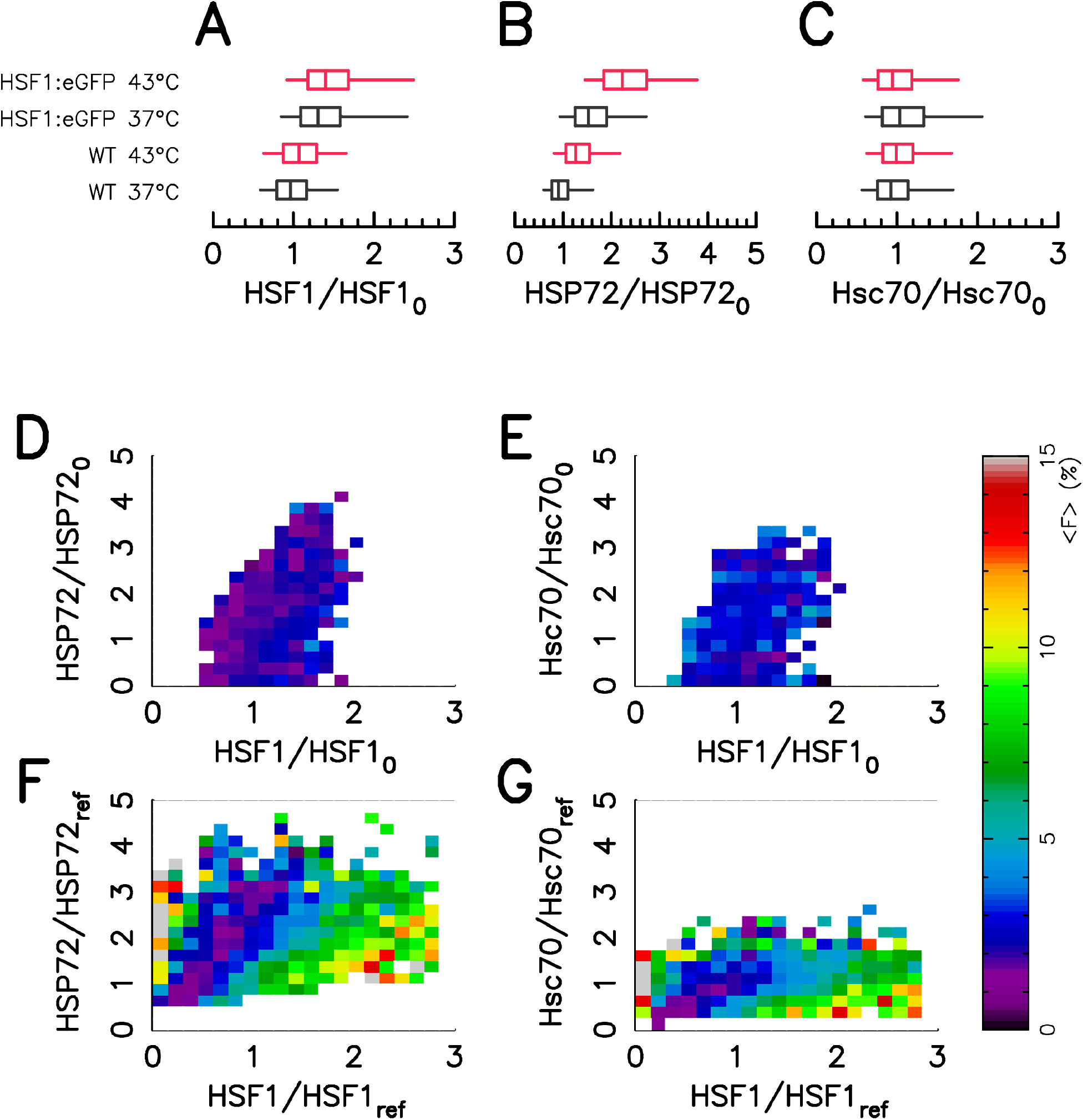
Foci intensity vary with protein concentration. A-C – Protein expression distribution measured by Immuno-Fluorescence in HeLa Wild Type and HeLa–HSF1:eGFP cell lines for two thermal condition: at a 37°C temperature (black), at a 43°C temperature for one hour (red). The box represent quartile and the whiskers represent the the 6th percentile and the 95st percentile. The distributions are displayed in Fig. SI 6 of the supporting information. D-E – Average value of foci intensity in HeLa Wild Type cell line for a given HSF1 and HSP72 level of expression (D) and for a given HSF1 and Hsc70 level of expression (E). F-G – Average value of foci intensity in HeLa–HSF1:eGFP cell line for a given HSF1 and HSP72 level of expression (F) and for a given HSF1 and Hsc70 level of expression (G).

In a second step we quantify the influence of HSP and HSF1 expression levels on the response amplitude. To do so in HeLa WT cells we estimate HSF1 and HSP72 (or HSC70) concentrations *via* double immunolabeling. One hour after exposure to a step temperature increase at 43°Cwe measure *F* for the whole cell population from HSF1 immunolabel. We then compute the population average of *F*(〈*F*〉) conditional to a given value of HSF1 and HSP72 immunofluorescence signals (Fig. 4 D). In agreement with the model prediction (Fig. 3) 〈*F*〉 increases with HSF1 level and decreases with HSP72 level. In contrast no significant correlation is found between 〈*F*〉 and HCS70 protein expression level (Fig. 4 E).

As shown above (Fig. 4 A-C) the HSF1:eGFP insertion increases the number of cells having higher concentration of HSF1 and HSP72. We perform HSP72 (or HSP70) immunolabeling in HeLa HSF1:eGFP one hour after exposure to a temperature step-up from 37°C to 43°C to compute maps similar as in (Fig. 4 D-E) with a stretched variability in proteins distribution (Fig. 4 F-G). In this case we monitor HSF1:eGFP fluorescence to measure HSF1 expression level and *F*. The stretched protein distribution makes the dependence of 〈*F*〉 on HSP72 protein expression level even more obvious. Immunolabelling of HSC70 reveals a dependence of 〈*F*〉 also on HSC70 concentration. Importantly we note that the average *F* value is similar in HeLa WT and in HeLa HSF1:eGFP experiments for a given HSF1–HSP couple. The mapping of *F* conditional to HSF1 and HSP concentration in HeLa HSF1:eGFP cells allows us to monitor rare events in which HSF1 and HSP72 concentration are higher.

We finally compute the percentage of the *F* dispersion explain by Eq. 8 by using InterQuartile Range (IQR) as a measurement of the data dispersion. We apply the procedure on data obtained with HeLa HSF1:eGFP cell line because the protein distributions are broader, and thus less sensitive to noise estimation of *F*. It turns out that Eq. 8 explain 52% of the dispersion by mapping *F* with HSF1 and HSP72 concentrations and 43.3% of the dispersion by mapping *F* with HSF1 and HSC70 (see Fig. SI 7 of the supplementary information). We note that the value of the exponent 3 (which reflects HSF1 trimerization) in Eq. 8 is crucial to explain the data dispersion (see SI 8 of the supplementary information). Indeed without it the percentage explaining *F* distribution falls to 41% with HSF1 and HSP72 and to 0% with HSF1 and HSC70.

## DISCUSSION

This study aims to better understand the molecular origin of phenotypic heterogeneity that will be crucial to work around resistance of a sub-population of cells upon cancer treatment. We also provide here a test for predictability of phenotypic heterogeneity by a mathematical model of the HSRN. We focus on HSRN because most anti-cancer treatments will activate this stress response. Moreover HSRN is a good model system as it is well characterized in the literature and heat shock provide a homogeneous way to treat an entire cell population.

We combine high-throughtput single-cell fluorescence experiments and mathematical modeling of the genetic regulation network. We first highlight an unexpected wide cell-to-cell variability in the activation of HSRN. We show that variability of the heat stress response is largely the result of cell–to–cell basal HSPs level heterogeneity. Surprisingly only a narrow variation in HSPs basal level is sufficient to induce the observed cell-to-cell variability. In the model, heterogeneity amplification is induced by the sequestration mechanism at the core of HSRN activation. Immunofluorence labeling experiments confirm that such an HSP expression level distribution explains a significant fraction of the heterogeneity and that the response amplitude depends on HSP expression level *via* the predicted mathematical relationship.

The response continuum observed in HSRN activation is a novel and surprising result that both completes the well–established biochemical data in the field and renew their interpretation. Owing to the fact that sequestration cascade leads to response hypersensitivity this sequestration cascade induces a heterogeneity amplification for amplitude stimuli close to the response threshold. At a fixed stimuli level the initial state of the cellular proteome determines the HSRN cellular activation response. This response may be mainly classified in three phenotypic clusters: (1) no activation, (2) transient activation, or (3) sustained activation. In a cell population all cells have a different proteome and thus all three types of activation dynamics are found for a given stress stimuli. We found that the probability of sustained activation increases with the stimuli level. Therefore, the averaged dynamical response measured by biochemical measurements (averages over a cell population) characterizes more the occurrence of the various phenotypes than the dynamics associated with a specific stress stimuli. Furthermore, although the sequestration cascades are heterogeneity amplifiers for chronic stress (T>=43°C) the same sequestration cascades induce heterogeneity collapse in the case of a mild stress. With this viewpoint hypersensitivity of the stimuli-to-response curve could be a strategy to quench the protein expression heterogeneity below a given stimuli threshold.

We show that variation of HSP72 and HSC70 molecular chaperones plays a major role in CCRV. This is expected within the framework of titration model for HSF1 as HSP70 family taht is shown to interact strongly with HSF1 transcription factor in unstressed cells Shi *et al*, 1998. This interaction was shown to be responsible for HSF1 sequestration in the absence of stress and desequestration from HSP70 that is to be crucial for HSRN activation in yeast Zheng *et al*, 2016. We have tested whether CCRV could also arise from level variations of HSP90, another important chaperone. While HSP90 exhibits weakly interaction with HSF1 without stress Zou *et al*, 1998, recent results suggest that sequestration may not be the important role of HSP90 in HSF1 regulation Kijima *et al*, 2018. Instead HSP90 interacts with transcriptionally active HSF1 trimers Conde *et al*, 2009 and newly synthetized HSP90 may regulate HSF1 by attenuating its ability to promote transcription when bound to HSE in DNA Kijima *et al*, 2018. Interestingly, at the single cell level, we do not find correlation between HSRN activation and HSP90 copy number (data not shown). These results are consistent with the fact that our readout (nSBs) is a measurement of HSRN activation but does not reflect transcriptional activity *per se*.

Human HSP70 expression was shown to vary with the cell cycle stage Milarski and Morimoto, 1986. However, in our experimental conditions, we do not find a significant correlation between single cell DNA content (assessed by Hoechst fluorescence level) and HSP72 or HCS70 nuclear concentration. Instead HSP expression level distribution could be attributed to transcriptional bursts intrinsically amplified by mRNA processing that causes substantial noise amplification at proteins level (Hansen and O Shea, 2016). Within our mathematical framework, HSP72 and HSC70 copy number explain around 50% of CCRV. Recent experiments in yeast have revealed that HSF1 hyperphosphorylation is another source of variability in HSRN Zheng *et al*, 2018. Such post-translational modifications control HSF1 activity on HSE rather than its activation in the cytosolZheng *et al*, 2016. We note that HSF1 phosphorylation could play a role in the CCRV we observe as it might induce variations in HSP72 transcription rate upon stress. However, it has to be noted that in our study we focus on activation of HSF1. Newly produced HSP72 (one hour after the stress onset) is rather small compared to pre-stimulus HSP72 level (see Fig. 4) and the span of HSP72 expression level is comparable to the one of unstressed cells (Fig. SI 6). Moreover the effect of HSC70 (which is not stress induced) on CCRV confirms the existence of a variability source, distinct from HSF1 phosphorylation, where the pre-stimulus cellular state at least partially determines single cell stress response.

HSP72 and HSC70 play a similar role in the refolding of misfolded proteins. Expression levels of the two protein subspecies are not correlated. From the network point of view this redundancy is intriguing. We suggest that such a redundancy may help to quench CCRV over the cell population. To test this hypothesis we include in the mathematical model two HSP species having uncorrelated but similar expression levels. This result in a reduced response heterogeneity with a two fold lower standard deviation (Fig. SI 9 of the supplementary information). HSPs redundancy may therefore reflects a strategy to compensate the protein expression fluctuations.

Our results highlight that the sequestration cascade mechanisms leading to the titration of HSF1 by basal HSP and MFP can control with ultrasensitivity the stress response. It is a sufficient guideline of a regulation network that describes the cellular heat shock response at both the population and the single cell levels. In this latter case, the HSPs stochastic expression variability explains the observed phenotypic heterogeneity. Therefore the ability to control the HSPs expression distribution, and not only its averaged expression level, should imply the ability to control the phenotypic heterogeneity and then to potentially reduce a therapy resistant subset of cells. Hypersensitivity of HSNR is a feature shared by several stress induced biological networks. As the amplification of heterogeneity is due more to hypersensitivity of the response than to the molecular mechanisms that engender it (sequestration in our case) the results and methods developed here could therefore be extended to other networks of stress and other hypersensitive networks.

## MATERIALS AND METHODS

### Cell culture and cell transfection

The HeLa human cervical cancer cell line (CCL-2™) was purchased from the American Type Culture Collection (ATCC, Manassas, VA). These adherent cells are grown as monolayer in Dulbecco’s modified Eagle’s medium (DMEM; Lonza, Levallois-Perret, France) supplemented with 10% (v/v) fetal bovine serum (FBS; Life Technologies, Saint-Aubin, France), 1% L-glutamine (2 mM) and 1% (v/v) penicillin-streptomycin (100 IU/ml) (Lonza). Cell cultures are maintained at 37°C in a humidified atmosphere containing 5 % CO_2_ (v/v), and passage at preconfluence (twice a week) using 0.05 % trypsin-0.53 mM ethylenediamine tetraacetate (EDTA; Lonza). HeLa growing cells are routinely screened for the presence of mycoplasma using DNA-staining with the nuclear dye Hoechst 33342 (1:10000 dilution) (Sigma-Aldrich, L’Isle d’Abeau Chesnes, France) to avoid collecting data from unknowingly contaminated cell cultures.

Wild-type HeLa cells (HeLa-WT) were transfected with a plasmid expressing the human full-length HSF1 fused to eGFP. The plasmid was kindly provided by Dr. Claire Vourc’h (Université Joseph-Fourier, Grenoble, France) and built as previously described Herbomel *et al* (2013). Briefly, PCR amplification allowed to obtain the coding sequence for human HSF1 that was cloned into peGFP N3 vector (Clontech Laboratories Mountain View, CA); the plasmid was then verified by sequencing (GATC Biotech, Constance, Germany). The transfection (of wild-type HeLa cells with the HSF1:eGFP plasmid) was carried out using FuGENEï¿œ HD transfection reagent (Promega, Charbonnières, France) according to the manufacturer’s instructions. The stable HSF1:eGFP-transfected (HeLa-HSF1:eGFP) cell line was then established under selective pressure by 1000 μg/ml geneticin (Life Technologies) followed by selectionf of a single GFP-positive cell by flow cell sorting system (FACSAria III, Becton Dickinson, San Jose, CA).

All experiments were performed on 2-day-old cell cultures (50 % confluence) prepared by seeding 1.8×10^5^ cells into 35-mm dishes (Sarstedt, Marnay, France) in complete DMEM without phenol red.

### Immunofluorescence staining of HSPs and HSF1

After 48 h of culture, HeLa-WT and HeLa-HSF1:GFP cells are heated at 43°C for one hour in our homemade incubator controlled in temperature and gas conditionsAnquez *et al*, 2012. At the end of the thermal stress, and in parallel to the unstressed samples (control, 37°C), cells are immediately rinsed with Dulbecco’s Phosphate-Buffered Saline (DPBS; Lonza), and fixed in 4% paraformaldehyde in DPBS for 10-15 minutes at room temperature (RT). After washing three-times with DPBS, samples are incubated for 30 minutes at RT in DPBS containing 0.3% Triton X100 and 5% goat serum (v/v) allowing permeabilization of cells and blocking of non-specific binding sites. Cells are then incubated overnight at 4°C with monoclonal primary antibody as following: mouse anti-Hsc70 (1:100 dilution; Santa Cruz Biotechnology, Dallas, Texas, USA), or mouse anti-HSP72 (1:100 dilution; Enzo Life Sciences, Villeurbanne, France), or rabbit anti-HSF1 antibody (Enzo Life Sciences). Subsequently, samples are washed, and incubated for 90 min at RT with either a goat anti-mouse (for Hsc70/HSP72 expression) or a goat anti-rabbit (for HSF1 expression) secondary antibody conjugated to Alexa Fluor-594 (Life Technologies). A DNA-staining using Hoechst 33342 (1:10000) is also performed for all samples, to allow the automatic detection of nuclear areas for image analysis.

### Live/Fixed cells imaging

HeLa cells are cultured in 35 mm dishes (starsedt) at approximately 50% confluence. Samples are placed on a Nikon TiE microscope with motorized filters wheel equipped with a XY-motorized stage (ASI). Cells were imaged through a 40X microscope objective (NA=0.6, Nikon) on a sCMOS camera (OrcaFlash LT, Hamamatsu). We set the camera binning to 2 resulting in an effective pixel size of 325 nm. Illumination for fluorescence and brightfield imaging is achieved through custom built optical system (components from Thorlabs). We use LED light source (Thorlabs) for synchronization of illumination with other apparatus. Exposure time is set to 150 ms for all experiments and for each fluorescence channel as well as brightfield illumination. Light power density, filter set and LED for each type of experiments are summarized in Tab. SI 1 of the supporting information. We use a custom-built acquisition software written in Labview to control the setup.

For live cells experiments culture dishes are maintained in a custom-built incubator which regulate temperature, humidity and atmosphere. The incubator was described inAnquez *et al*, 2012. Cells are maintained at 37°C for one hour and heat shocked at 41, 42 or 43°C for three hours by increasing the incubator temperature. Time evolution of the Nuclear Stress Bodies foci is monitored in real time. In order to increase the output rate of the experiment we acquire data for ten different fields of view in the same dish (by use of the motorized stage) leading to the tracking of approximately 200 cells per experiment. Two consecutive fields of view are separated by approximately 300*μ*m. To account for focusing drift and to allow image segmentation we acquired for each filed of view a z-stack of nine images per channel by moving the objective lens along the optical axis. Two consecutive images of the stack are defocused by approximately 2*μ*m. We acquired z-stack at a 0.5 image/min rate.

For fixed cells imaging heat shock are performed in the same incubator as the live cell data. Cells are shocked for one hour at 43°C and then fixed right after. After fixation and immunolabeling (protocols described below) cells are imaged on the same microscope. For these experiments we used the Nikon Perfect Focus system and thus did not acquire z-stack. We image 400 positions per condition leading to approximately 8000 cells per experiment.

### Image processing and analysis

All image processing and data analysis was performed using custom written algorithms either in Fortran or in Matlab.

For time lapse microscopy experiments we first estimate best focus for each z-stack by use of a contrast function Price and Gough, 1994. Best focus was estimated from fluorescence images. We did not found significant defocusing between fluorescence and brightfield in our experimental conditions. Cells were automatically segmented using brightfield image z-stack. For this we take advantage of the fact that gray level varies across the z-stack for pixels located in cells while such gray level is approximately constant for background pixels and pixels at the periphery of the cells. Image segmentation was visually inspected after image processing. Corrections to cell segmentation were carried when necessary *via* a custom written semi-automated graphical interface by either removing false positive or correcting masks. After cell segmentation cells were tracked by simply linking the closest cell found in the next image. Visual inspection of the tracking did not reveal errors as the cells do not move significantly in the time interval between two acquisition. We then estimate background for fluorescence images by convolving the raw data with a 30 pixels wide gaussian kernel (larger than cell size) and averaging across the z-stack. Background was subtracted to raw data for further analysis. Total HSF1-GFP intensity was simply estimated by integrating fluorescence intensity over the whole cell mask. HSF1-GFP foci were automatically detected by use of wavelets transform with wavelet radius of 2 pixels Olivo-Marin, 2002. Only spots with maximum intensity higher than mean cell intensity was considered for further analysis. The *F* factor was defined as the integrated intensity found in all foci divided by the total cellular fluorescence.

For fixed cells immunofluorescence experiments image segmentation was achieved on images from HSP fluorescence channel for whole cell segmentation and on the images from Hoechst fluorescence channel for the nucleus segmentation. We acquired fluorescence images of dishes filled with fluorescent dye for flat field correction. The dyes were courmarin for Hoechst channel, rhodamine 110 for GFP and AlexaFluor488 channel and rhodamine B for AlexaFluor594 channel. After flat field correction images were segmented using a modified Otsu thresholding method Otsu, 1979. A constant background was subtracted before further analysis. *F* was defined as above and HSP concentration was defined as the total fluorescence inside nucleus divided by the nuclear area in arbitrary units. False positive detection were removed by selecting a polygon in the Hoechst intensity versus nucleus area plane.

### Mathematical model for HSRN

The heat stress cellular response dynamic is mainly regulated by two complexes that both involve the chaperone proteins HSP (Sivéry *et al*, 2016). HSPs titrate the misfolded proteins, on the one hand, and its own transcription factor (HSF1) on the other. A reduced model of the cellular response to heat stress is constructed from a detailed kinetic one of the literature (Sivéry *et al*, 2016) under the following assumptions: (i) all protein species have similar half-life; (ii) the assembly dynamics and assemblies of the protein complexes are adiabatically eliminated and equilibrium equations at the fixed points are approximated by rational functions (See Sec. SI 11 of the Supporting Information for details).

In addition to the model developed in Sivéry *et al*, 2016, the present model improves the regulation of the translation process via HSPs. In fact, Heat shock proteins are requested to initiate the translation process, therefore the sequestration of Heat shock Protein by Misfolded Protein reduces the ability of HSP to initiate the translation. Straightforwardly, we include in the modeling an HSP dependent translation rate which decreases as free monomeric HSP form vanishes. This mechanistic detail is crucial to describe the slow increase of foci dynamics during a 43°C heat stress.

The model equations reads:

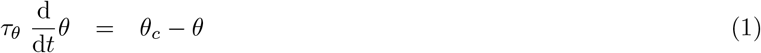

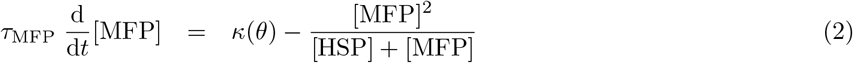

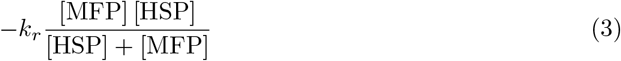

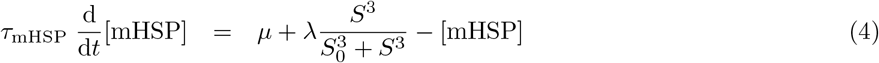

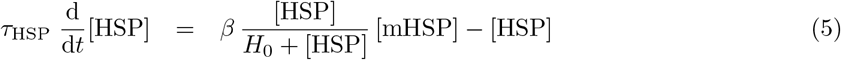

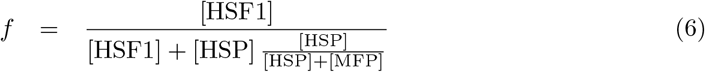

where *t* is time; *θ* is the temperature of the cell environment measured in °C; [MFP], the misfolded protein concentration; [mHSP], the concentration of mRNA coding for HSP; [HSP], the heat shock protein concentration; [HSF1] the heat shock factor 1 protein concentration; and *f*, the concentration of free HSF1 proteins (not bounded to HSP).

The denaturation rate *κ*(*θ*) is here the only temperature input. Mathematical expression of *κ*(*θ*) was discussed in Peper *et al*, 1998 and takes the following form in the range 37°C–45°C:

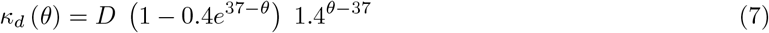

Note that *τ_θ_*, the incubator rise time, was measured experimentally. Also all rate constant are normalized (see Supporting information for details). The signification and values of model parameters are summarized in Tab. SI 2 of the Supplementary Information.

In this framework, the fraction of HSF1 bound to NSBs at a given date *t* is proportional to:

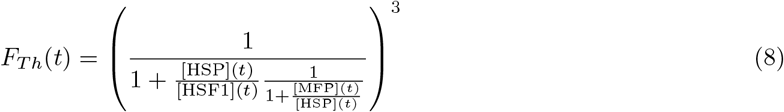

where [HSF1](*t*), [HSP](*t*), and [MFP](*t*) refer to the HSF1, HSP, and MFP concentrations at the date *t* (Fig. 3-A). Note that the power 3 in expression Eq. 8 arises from the fact that only trimmers of HSF1 are bound.

This expression for *F_Th_* given in Eq. 8 depends on two concentration ratios, 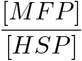 and 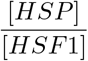 that reflect the two competitive complex formations: The first ratio reflects the MFP titration by HSPs while the second ratio accounts for HSF1 titration by HSPs.

In practice we seek to compare measurements from fluorescence microscopy with expression 8. Concentration of HSF1 and HSP are measured in fluorescence (arbitrary) units while MFP concentration is unknown. We introduce a scale parameter *α* that both account for conversion from fluorescence units to concentration units and the unknown concentration of MFP relative to HSP. It is important to note that *α* can vary from one cell to another only because MFP concentration is unknown. For convenience we introduce 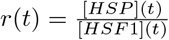 where [*HSP*](*t*) and [*HSF*1](*t*) are measured in fluorescence units. *F_Th_*(*t*) then reads:

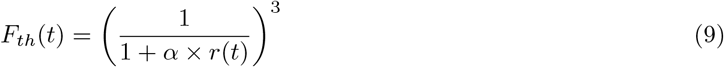

## Acknowledgements

This work has been partially supported by the LABEX CEMPI (ANR-11-LABX-0007), as well as by the Ministry of Higher Education and Research, Hauts de France council and European Regional Development Fund (ERDF) through the Contrat de Projets Etat-Region (CPER Photonics for Society P4S).

## Author Contributions

Performed research M.G. F.A. A.P. Q.T.; Designed research M.G. F.A. E.C. Q.T.; Analyzed data; M.G. F. A. Q.T.; Wrote the paper F.A. E.C. Q.T.

## Supplementary Information

### SI 1 Imaging conditions

**Table SI 1:** Imaging conditions

| Fluorophore | Light source | Excitation | Dichroic | Emission | Light |
| --- | --- | --- | --- | --- | --- |
| - | LED | filter | mirror | filter | intensity |
| - | (Thorlabs) | (Semrock) | (Semrock) | (Semrock) | ($W.cm^{-2}$) |
| GFP | M490L4 | FF02-482/18 | FF495-Di03 | FF02-520/28 | $\sim 1.8$ |
| AlexaFluor488 | M490L4 | FF02-482/18 | FF495-Di03 | FF02-520/28 | $\sim 1.8$ |
| AlexaFluor594 | MCWHL5 | FF01-536/20 | Di02-561 | FF01-600/37 | $\sim 1.35$ |
| Hoechst | M365LP | FF01-360/2 | FF416-Di01 | FF02-460/80 | $\sim 0.63$ |
| Brightfield | MCWHL3 | - | FF495-Di03 | FF02-520/28 | $\sim 2.25$ |

### SI 2 Foci intensity dynamic in wild-type cell line

**Figure SI 1:**
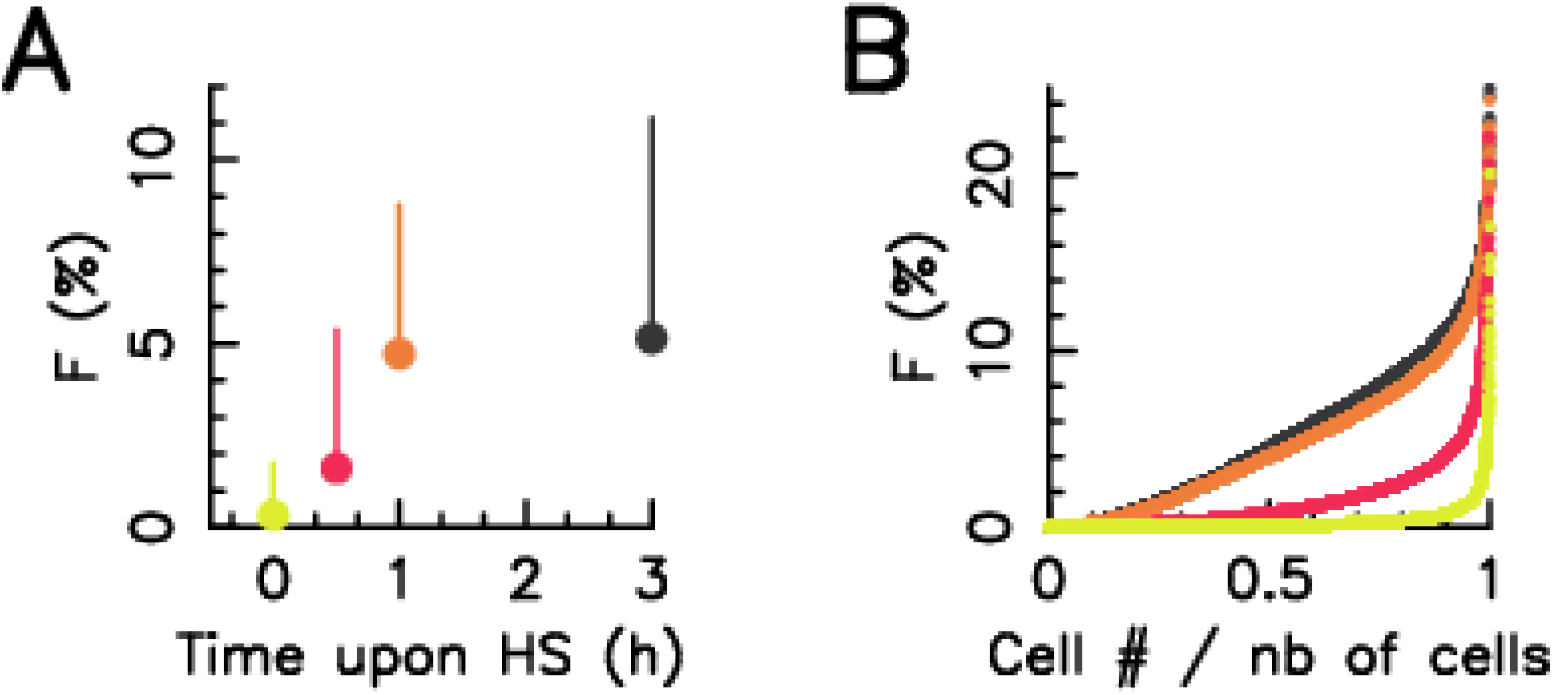
Foci intensity dynamic in wild-type cell line during a 43°C Heat stress. A – Averaged value over the cell population of foci intensity measured on single-cell (dots) and associate error bar. B – Foci intensity in single cell ordered by increasing value. In both pictures the color code is the following: yellow −37°C; red - half an hour after the oncet of a 43°C; orange - one hour after the onset of a 43°C; black – three hour after the onset of a 43°C.

### SI 3 Total HSF1:egfp expression level do not vary during experiments

**Figure SI 2:**
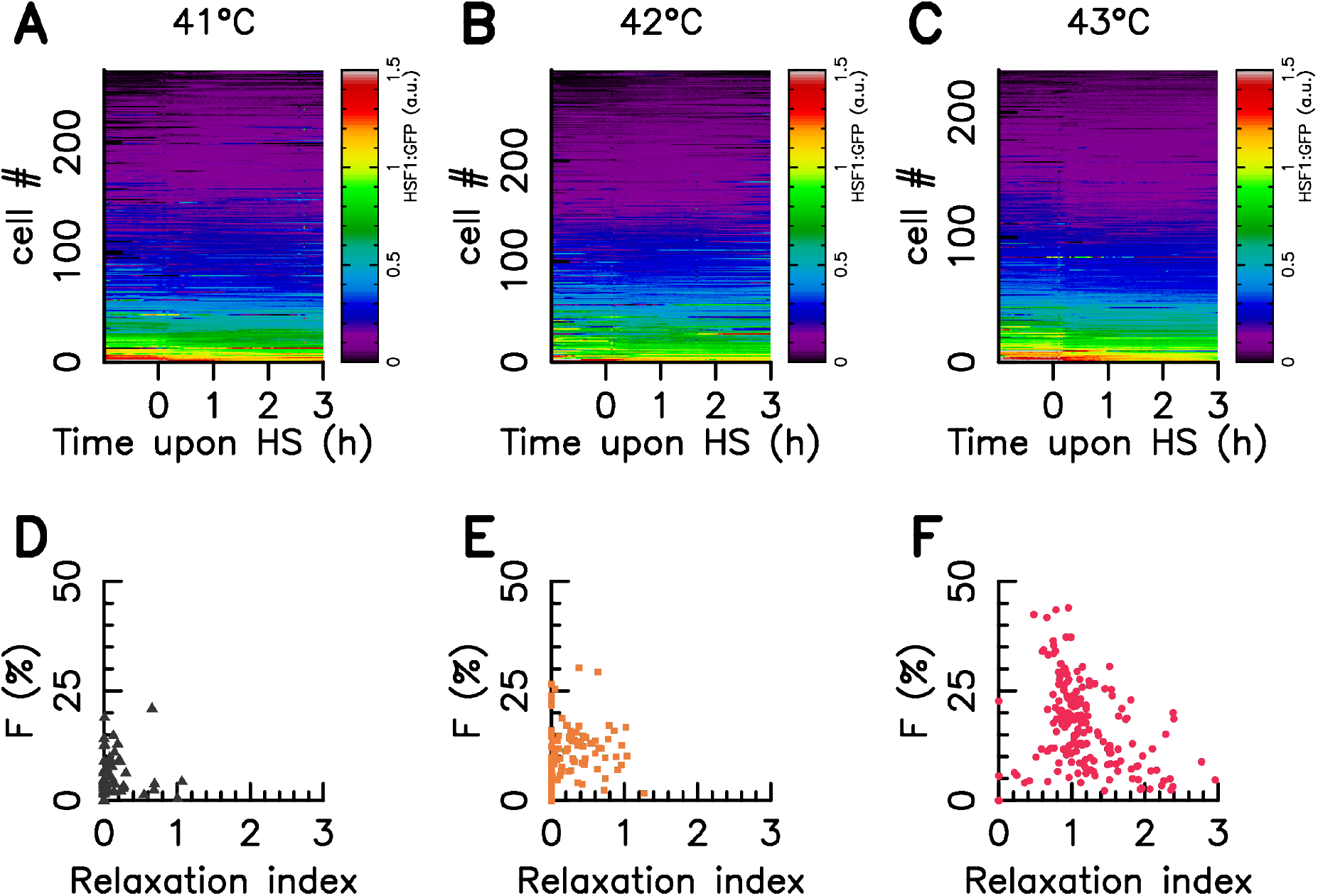
Additional results of Figure 2. A-C HSF1:eGFP fluorescence is monitored over time in single cell upon a 41°C 42°C and 43°C heat stress, time zero coincides with the stress onset. Each horizontal line corresponds to a single cell, a color code indicates the HSF1 fluorescence measured at a given time, according to the scale bar on the right. D-F Correlation between Fraction of HSF1 fluorescence within foci one hour after the stress onset and the relaxation index defined as the ratio of the foci intensity measure in a given cell at three hours after the stress onset to the one measured one hour after the stress onset.

### SI 4 High–troughput screening of Foci Dynamics with the use of another HeLa cell line transfected with the same HSF1– eGFP construct

We check the influence of the transfection on the experimental result The experimental result display in Fig. 2 are confirmed by the use of another HeLa cell line transfected with the same HSF1–eGFP construct.

**Figure SI 3:**
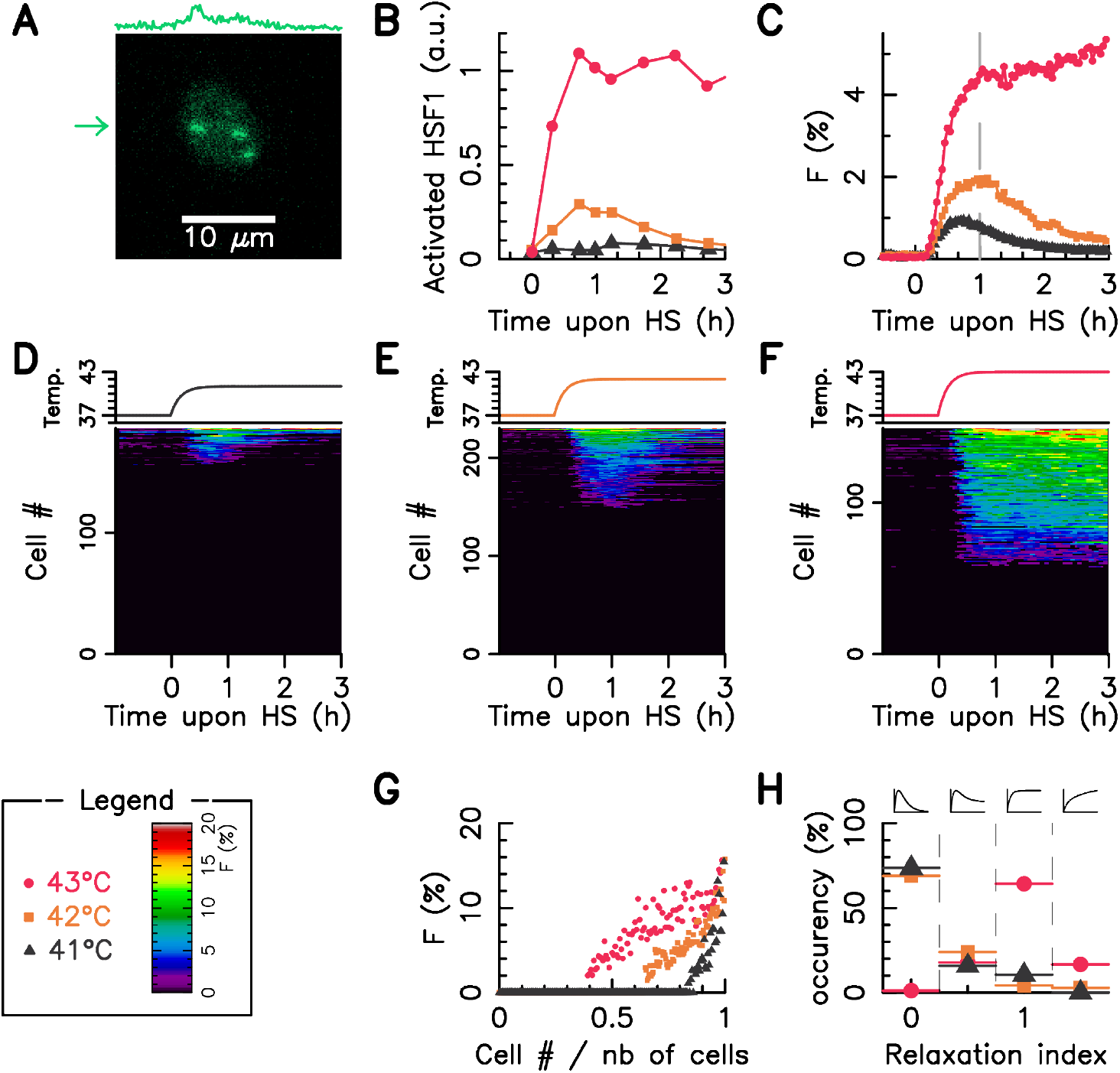
High—troughput screening of Foci Dynamics in individual HeLa cells. Result obtained with another HeLa cell line transfected with HSF1:eGFP construct. The faction of HSF1-eGFP fluorescence is monitored over time in single cell upon a 41°C, 42°C, and 43°C heat stress, time zero coincides withe the stress onset. A – A single HeLa HSF1:GFP cell, 1 hour after the onset of the 43°C Heat stress with three visible Foci; the profile of intensity for the row indicated by the arrow is displayed on the top of the box. B – Dynamics of activated HSF1 as measured as measured by Abravaya *et al.*. C – Average value over the cell population of foci intensity. D-F Cell temperature time profile (upper panel) and fraction of HSF1 fluorescence within foci in single cell (lower panel); in the color image, each horizontal line corresponds to a single-cell, a color code indicate the faction of HSF1 fluorescence within foci measured at a given time. G – Fraction of HSF1 fluorescence within foci variation over the cell population one hour after the stress onset (cell ranking is similar to D-F). H – Statistical distribution of the relaxation index defined as the ratio of the foci intensity measure in a given cell at three hours after the stress onset to the one measured one hour after the stress onset. The legend box defines the used color code.

### SI 5 Model Parameter Estimation

**Figure SI 4:**
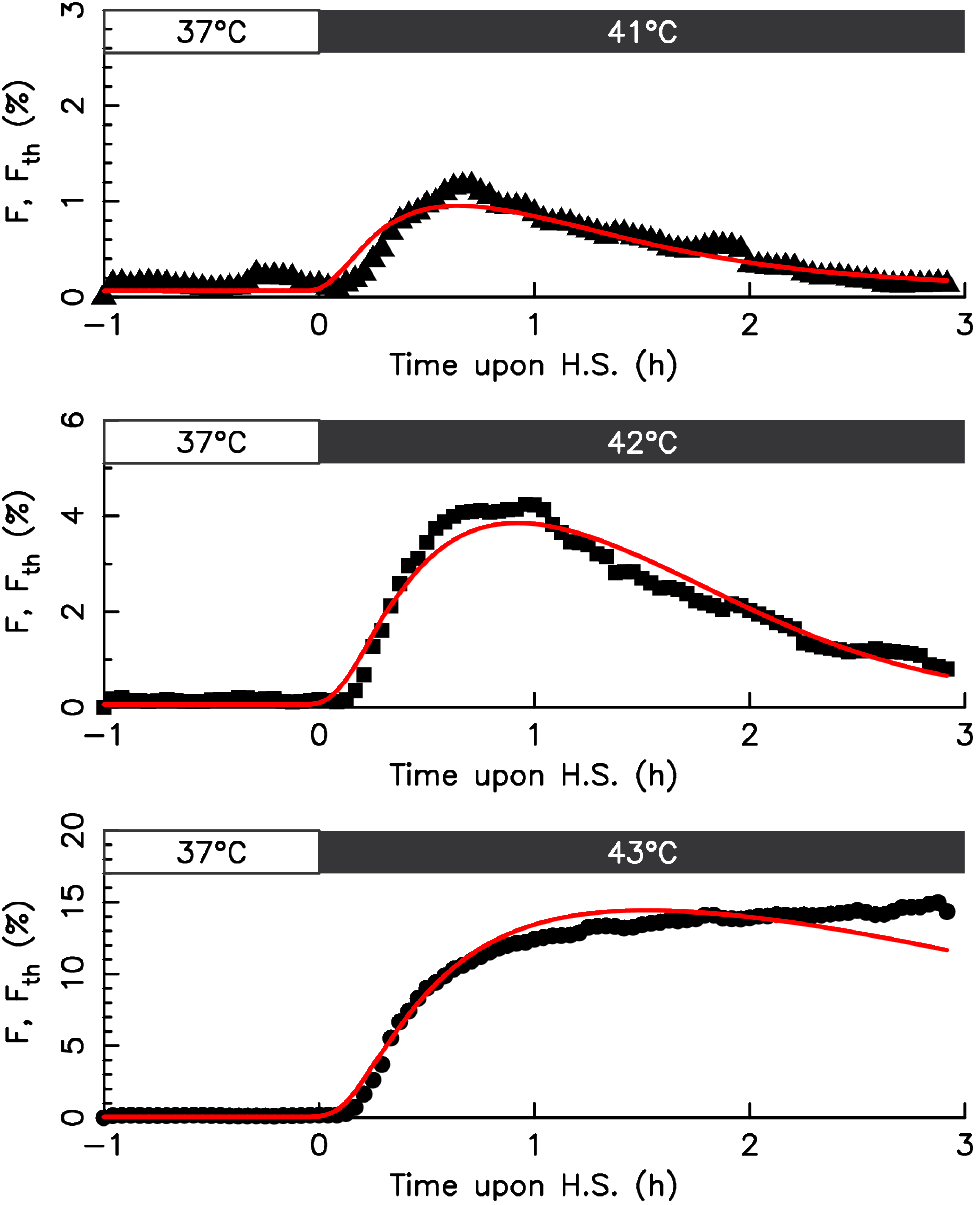
Best adjustment of average foci value by the reduced model. Model output (red lines) and mean foci value (points). Time zero coincides with stress onset, the thermal protocol is indicated by bars on the top of each boxes. The root mean square deviation value 0.1 % for 41°C, 0.26 % for 42°C, 1.5 % for 43°C, and 0.9 % alltogether.

**Table SI 2:** Estimated Parameter of the heat shock response network

| Parameter | unit | description | value |
| --- | --- | --- | --- |
| $k_D$ | ($\mu\text{M}$) | denaturation rate | 1.76 |
| $k_r$ | | renaturation rate | 17.7 |
| $\mu$ | ($\mu\text{M}$) | HSP basal transcription rate | $1.47 \cdot 10^{-3}$ |
| $\lambda$ | ($\mu\text{M}$) | HSP active transcription rate | 0.78 |
| $S_0$ | ($\mu\text{M}$) | HSP transcription regulation threshold | 0.18 |
| $\beta$ | | HSP translation rate | 10 |
| $H_0$ | ($\mu\text{M}$) | translation regulation threshold | 0.32 |
| [HSF1] | ($\mu\text{M}$) | HSF1 concentration | $4.0 \cdot 10^{-2}$ |
| $\tau_{Temp}$ | (h) | incubator rise time | 1/15. |
| $\tau_{MFP}$ | (h) | MFP lifetime | 0.5 |
| $\tau_{mHSP}$ | (h) | mHSP lifetime | 1 |
| $\tau_{HSP}$ | (h) | HSP lifetime time | 10 |

### SI 6 Coarse-grain model of Foci Dynamics and variability – Additional results to Fig. 3

**Figure SI 5:**
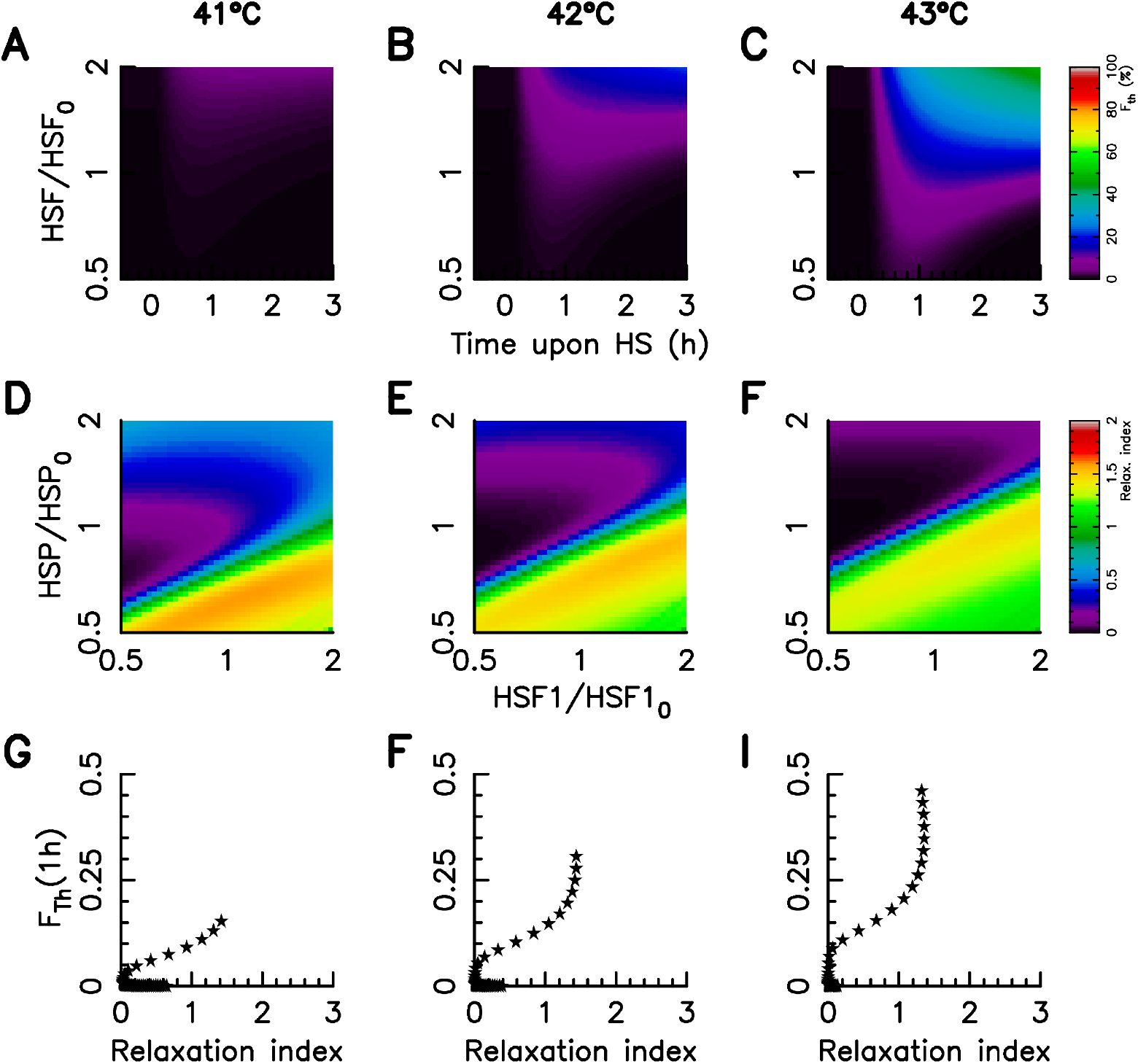
Coarse-grain model of Foci Dynamics and variability – Additional results to Fig. 3. A–C Relaxation index value for varying HSP and HSF1 copy number in the case of a 41°C (A), 42°C (B), and 43°C (C) heat stress. D-F Correlation between Foci intensity one hour after the stress onset and the relaxation index upon a variation of HSP.

### SI 7 Chaperon protein HSP72 and HSC70 are log-normally distributed

A random variable *X* follows a lognormal distribution of parameter (*μ*, *σ*) if it’s probability density function reads

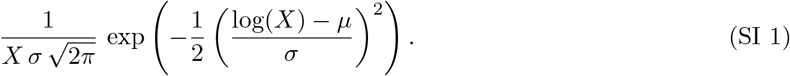

The mean reads E(*X*) = *e*^*μ*+*σ*^2^/2^.

### SI 8 IQR of the foci level variability description by HSP and HSF1

### SI 9 Foci correlates with *HSP/HSF1*

### SI 10 HSP redundancy may be a strategy to reduced the cell-to-cell variability

The cell-to-cell variability of the foci response arise from two fact: (1) The sequestration mechanism underlying the heat stress response induce a high sensitivity to protein expression level (Fig. SI 9-A) (2) The stochastic expression of genes induce protein expression level distribution (Fig. SI 9-A). The heat shock protein 70 family contains numerous homologous chaperone proteins (at least height) and HSP72 (the stress inductive chaperone) as well as HSC70 (the constitutive chaperone) which are the major members of the familly are log-normally distributed with a similar variance.

A question arise whether the HSP redundancy may be a strategy to reduced the cell-to-cell variability. To illustrate the phenomena, let us focus on the foci intensity one hour after a 43°C stress onset and compare two different cases. In the first case, the total pool of chaperon HSP is assumed to be build from a single gene and to follow a lognormal distribution of parameter (*μ*, *σ*^2^) (grey shade area in Fig. SI 9). In the second case, the total pool of chaperon HSP is assumed to be build from two distinct genes and then to be the sum of two homologous independent proteins, each of them following a lognormal distribution of parameter (*μ*′,*σ*^2^) and (*μ*′,*σ*^2^) (red lines in Fig. SI 9). In both case, the parameters *μ* and *μ*′ of the distribution are adjusted such as the mean of the total pool is unity, and the parameter *σ* is the same in all distributions. The duplication of chaperones in case two reduces by more than two the standard deviation of the foci intensity one hour after the stress onset.

### SI 11 Model reduction

#### SI 1 Adiabatic elimination of dimer assembly and dis-assembly

Let A and B be two proteins, the equilibrium equation of reversible complex formation reaction A + B↔A:B is written [A:B] *k*_0_ = [A] × [B] where *k*_0_ is a balance concentration. Straightforwardly, one gets

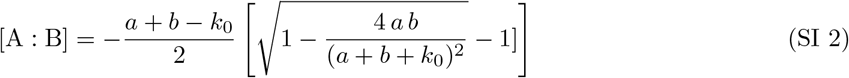

where *a* = [A] + [A:B] and *b* = [B] + [A:B] stands for the total concentration of proteins specie A and B. If now the chemical species concentration *a* and *b* dominates the equilibrium concentration k0 (*a* + *b* ≫ *k*_0_), parameter free rational functions approximate the concentration at equilibrium:

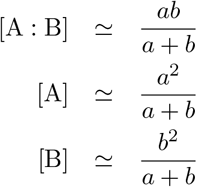

To reduce the mathematical model of the cellular heat shock response network, we consider that the hetero-dimer assembly and disassembly follow the equilibrium relation at equilibrium for a given pool of MFP, HSP, and HSF1, in a ordered reaction chain: firstly HSP binds MFP and secondly HSF1 bind the remaining free HSP pool. Let us denote by [*HSF*1]_*T*_, [*HSP*]_*T*_, and [*MFP*]_*T*_ the total concentration of HSF1, HSP, and MFP then the conservation relations reads

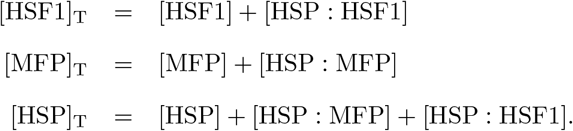

Once applied to the dominant hetero dimer reaction *HSP* + *MFP* → *MFP*: *HSP* the adiabatic elimination gives

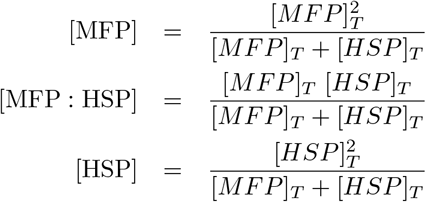

for the concentration of misfolded protein in free from, the hetero dimer MFP:HSP, and HSP in free form before HSF1 binding. In a second time, we compute the equilibrium between HSF1 and the remaining HSP in free form which leads to the expressions

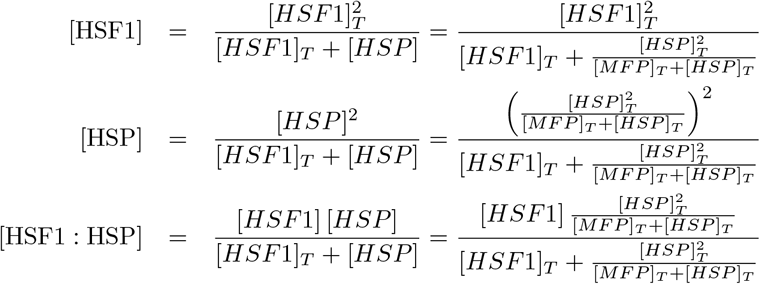

In the reduce dynamical model 1, only [HSF1]_T_, [HSP]_T_, or [MFP]_T_ appear as a protein concentration, the *T* subscript is thus removed for clarity.

**Figure SI 6:**
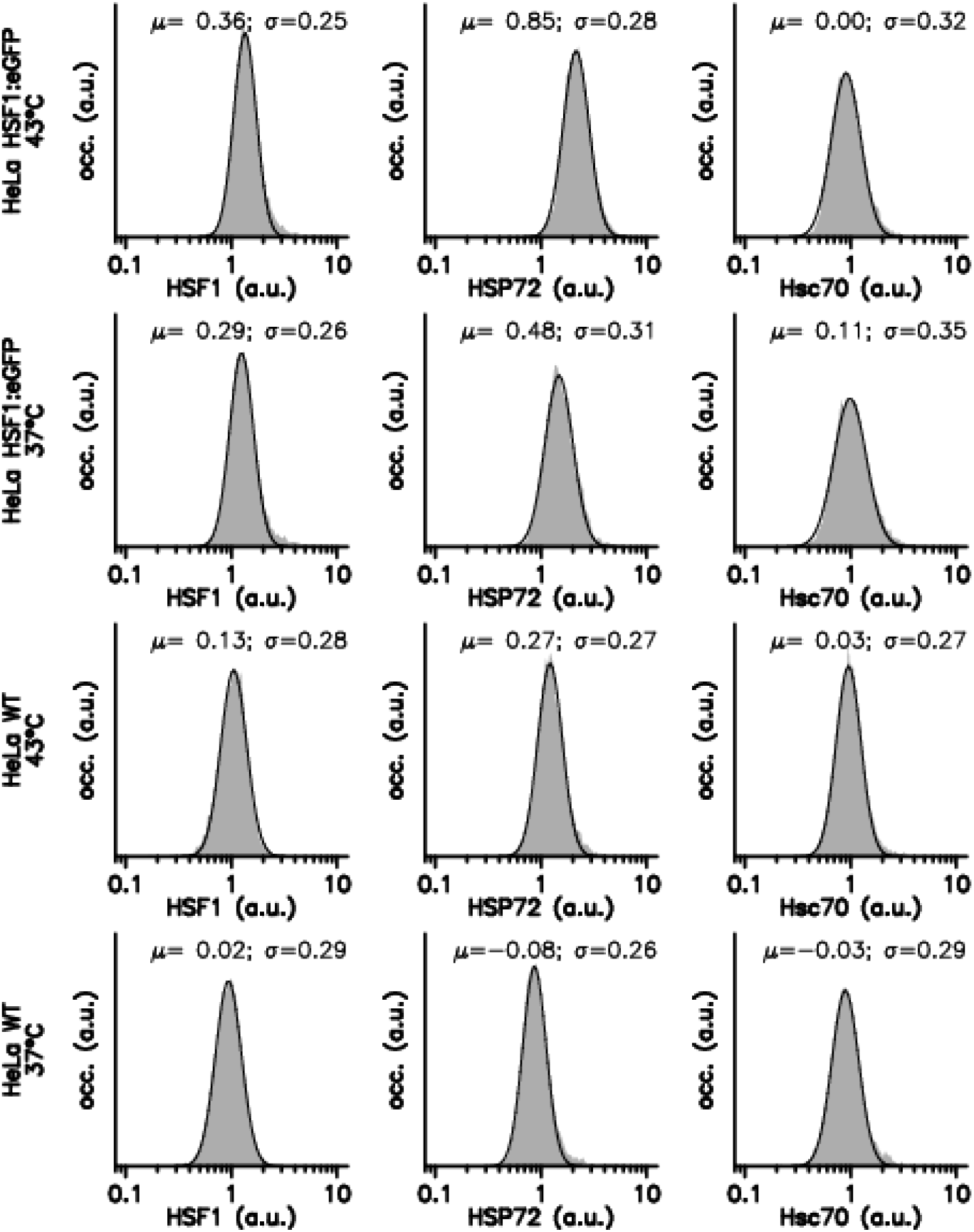
Protein level distributions. Protein expression distributions (grey shaded) are measured by Immuno-Fluorescence in HeLa Wild Type and HeLa–HSF1:eGFP cell lines for two thermal condition: at a 37°C temperature (black), at a 43°C temperature for one hour. The solid black lines correpond to best fitted lognormal distribution SI 1, estimated parameter values are written on the plot. Each line corresponds to a specific cell line and a specific thermal condition. Each column corresponds to a specific protein, HSF1, HSP72, and Hsc70. The distributions are normalised such that the mean in HeLa Wild Type at 37°C is unity.

**Figure SI 7:**
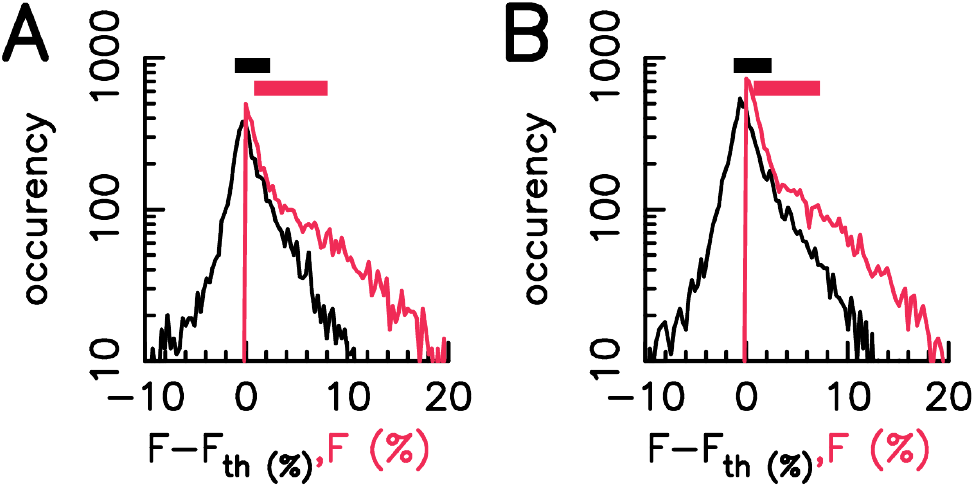
Black lines are histogram of the residuals between *F* and *F*_Th_ given by Eq. 8 using HSF1 and HSP72 (A) or HSF1 and HSC70 (B). Red lines are the histograms of *F* values. The filled square correspond to the IQR of the residual (black) or *F* value (red).

**Figure SI 8:**
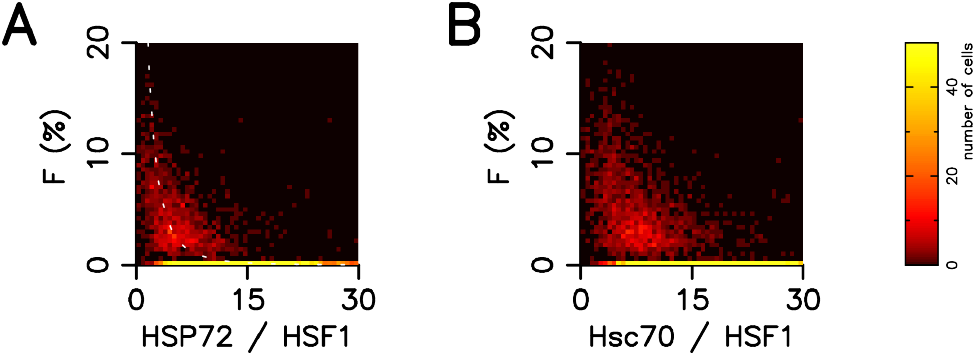
Cells distribution in HeLa–HSF1:eGFP as a function of Foci intensity (after 1 hour at 43°C) and HSP72 to HSF1 ratio (A) or Hsc70 to HSF1 ratio (B). In (A), the white dot line correspond to the mathematical function *y* = 1/(1 + *αx*)^3^ with *α* = 0.45

**Figure SI 9:**
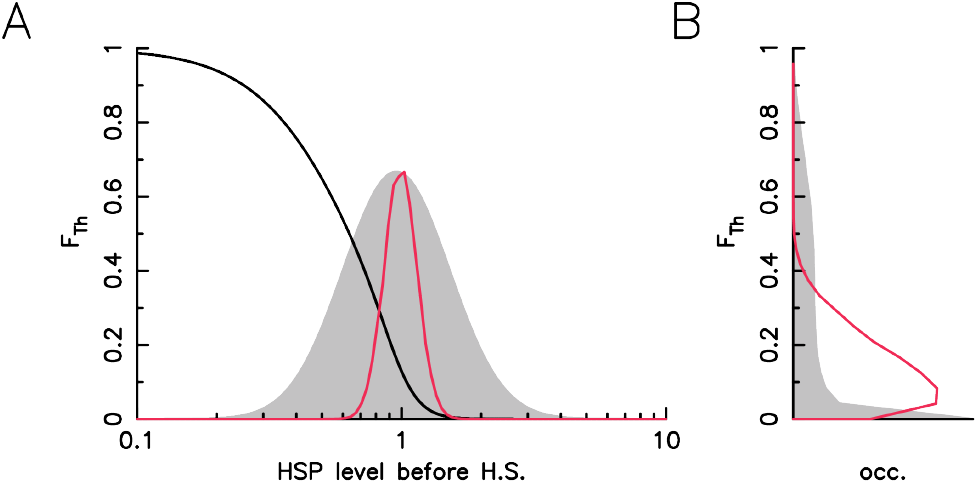
Protein redundancy quench the foci variability. A – Foci intensity one hour after a 43°C heat stress onset as a function of initial HSP in the mathematical model (black line), HSP distribution before the heat stress in the case of one HSP species log-normally distributed (grey shade area) or two uncorrelated HSP species log-normally distributed (red line). B - Corresponding foci intensity distribution for one HSP species (grey shade area) and two HSP species (red line).

